# Seed Microbiome Transfer Mitigates Intergenerational Dysbiosis, Modulates Plant Defenses and Suppresses Foliar Disease

**DOI:** 10.64898/2026.08.31.747941

**Authors:** Fabiano Jose Perina, Vanessa Thomas, Toi Ketehouli, Samerika Mudiyanselage, Mukesh Jain, Ina Schlathoelter, Erica Goss, Samuel Julio Martins

## Abstract

Antibiotic-induced disruption of plant-associated microbiomes has the potential to alter host health beyond the directly exposed generation, yet whether the effects of dysbiosis are transmitted through the seed microbiome remains unknown. Here, we investigated the intergenerational impacts of streptomycin-induced dysbiosis in tomato (*Solanum lycopersicum*), demonstrated that seed microbiome transfer (SMT) restores progeny microbiome function and disease resistance, and characterized the underlying physiological and genetic mechanisms. Parental streptomycin exposure altered the composition of progeny rhizosphere bacterial communities, reduced expression of defense-associated genes, and increased susceptibility to *Xanthomonas perforans*. Suppression of immune gene expression was strongly associated with increased disease severity, indicating that parental dysbiosis impaired progeny plants’ ability to mount effective immune responses. Transfer of the seed microbiome from healthy plant donors partially restored rhizosphere community composition, reduced disease severity and recovered defense gene expression of three genes. Together, our findings demonstrated that antibiotic exposure microbiome disturbance generates intergenerational legacy effects that influence plant immunity and disease susceptibility and seed microbiome transfer can counteract this dysbiosis across generations.

## Introduction

Plant-associated microbial communities are increasingly recognized as an extension of host functional capacity, contributing to nutrient acquisition, immune regulation, and protection against pathogens [1, 2]. The root adjacent soil, known as the rhizosphere, harbors the most diverse and extensively studied microbial community, followed by the phyllosphere and the seed microbiome [3–5]. Soil microbial communities associated with plant roots regulate key processes essential for plant and soil health, including nutrient cycling, organic matter decomposition, pathogen suppression, and immune priming, vital for plant and soil health [6–9]. Consequently, disruption of these microbial networks can have cascading effects on soil fertility, plant health, disease resistance, and long-term agricultural sustainability [10].

Agricultural management practices are major drivers of rhizosphere microbiome assembly, and broad-spectrum interventions can disrupt non-target microorganisms, resulting in unintended ecological consequences. Streptomycin, widely used to control bacterial plant diseases, can disrupt non-target soil microorganisms that perform essential ecological functions, while selecting for antibiotic-resistant microorganisms and resistance genes [6, 11, 12]. Such disturbances can lead to dysbiosis, broadly defined as a disruption in the composition, function, and stability of microbial communities that are associated with altered host or ecosystem health [13, 14]. In plants, dysbiosis has been linked to both biotic and abiotic stressors, impairing beneficial functions such as nutrient acquisition, pathogen suppression, and immune regulation [14]. For example, antibiotic-induced dysbiosis in the rhizosphere of tomato plants led plants to become more susceptible to foliar infection by *Xanthomonas perforans,* demonstrating that antibiotic-induced dysbiosis can extend throughout the plant [15]. Because *Xanthomonas* pathogens can also be seedborne, pathogen-mediated disruption of the plant seed microbiome may have implications for microbial transmission to the next generation. Additionally, disrupting the plant microbiome has been shown to negatively affect the seed microbiome [16], potentially increasing seed susceptibility to stress upon planting.

Rhizosphere dysbiosis can be mitigated through several approaches, including successive cultivation via the host’s *’cry for help*’ strategy [17] and through seed or rhizosphere microbiome transfer [18], which introduces microbial communities from healthy donors into dysbiotic systems to re-establish eubiotic states. In some cases, when missing host functions are identified, targeted microbial inoculation with the lost trait is sufficient to mitigate dysbiosis [19]. For example, Targeted inoculation with beneficial cyanobacteria reduced disease severity by restoring key ecological functions, illustrating a “patching the leak” approach without reconstructing the entire community [19]. Together, these strategies emphasize the central role of the microbiome in plant health, where microbial structure and function contribute to the extended plant phenotype [15]. Compared with pathogen-challenged plants harboring a eubiotic rhizosphere microbiome, plants with streptomycin-induced dysbiosis exhibited transcriptional reprogramming, including differential expression of *FLS2* (flagellin-sensing receptor), *CHI14* (chitinase), and *GluB* (*β*-1,3-glucanase), which was associated with greater disease severity [17]. Therefore, rhizosphere microbial communities can prime plant immunity by activating these and other defense pathways, increasing host readiness against pathogen invasion [1, 20, 21].

The persistence of microbiome-mediated effects on plant immunity across generations represents an emerging frontier in plant microbiome research [22, 23]. Previous studies have shown that parental environmental conditions and disease status can shape microbiome composition and influence subsequent plant generations [23, 24]. However, how antibiotic-induced dysbiosis alters microbial assembly and host functions beyond the exposed generation remains unclear. This represents a critical gap in understanding the long-term consequences of dysbiosis within the hologenome framework, in which host health and phenotype are shaped by interactions with associated microbial communities [25]. Microbiome transfer offers a strategy to restore functions lost during dysbiosis, yet whether it can reverse intergenerational consequences remains unexplored.

This study investigated rhizosphere microbiome composition, plant immune gene expression of selected genes, and disease severity in progeny plants to evaluate the intergenerational consequences of parental streptomycin-induced dysbiosis and to determine whether seed microbiome transfer (SMT) could restore protective rhizosphere functions. We hypothesized that parental streptomycin exposure would reduce rhizosphere microbial diversity in progeny, disrupt taxa associated with plant defense, and suppress immune-related genes linked to induced systemic resistance. We further hypothesized that SMT from healthy donor seeds would partially restore microbial community structure, enhance immune gene expression, and reduce disease severity caused by *X. perforans.* The objectives of this study were to (i) characterize the intergenerational effects of parental streptomycin exposure on the rhizosphere microbial community composition of progeny plants, (ii) determine whether seed microbiome transfer can mitigate antibiotic-induced dysbiosis and recover disease-suppressive functions within the rhizosphere, and (iii) evaluate changes in selected plant immune gene expression and disease progression following microbiome disruption.

## Materials and methods

### Experimental design and seed microbiome transfer and pathogen inoculation

To assess whether seed microbiome restoration mitigated the intergenerational effects of streptomycin-induced dysbiosis, seeds were collected from parental plants treated with sterile distilled water or streptomycin (0.6 g L⁻¹). Treatments were applied at 21 days after emergence and full anthesis as a 200 mL soil drench combined with foliar application to runoff. The parental experiment comprised two independent runs in a randomized complete block design (4 blocks; 3 plants per experimental unit). Seeds from 12 plants per treatment were pooled within each run, with seed lots from parental runs 1 and 2 used in progeny/SMT runs 1 and 2, respectively.

Progeny seeds were evaluated in a 2 × 2 factorial design crossing parental history (water or streptomycin) with SMT (±), with all seeds challenged with streptomycin-resistant *Xanthomonas perforans* strain XP1-6 before sowing. This generated four treatments: *Xp*+Water, *Xp*+SMT, *Xp*+Strep, and *Xp*+Strep+SMT (Figure 1). Two independent runs were conducted using a randomized complete block design (9 blocks; 3 plants per experimental unit; n = 27 per treatment). No run × treatment interactions were detected (*p* > 0.05); therefore, runs were combined for analysis.

**Figure 1.**
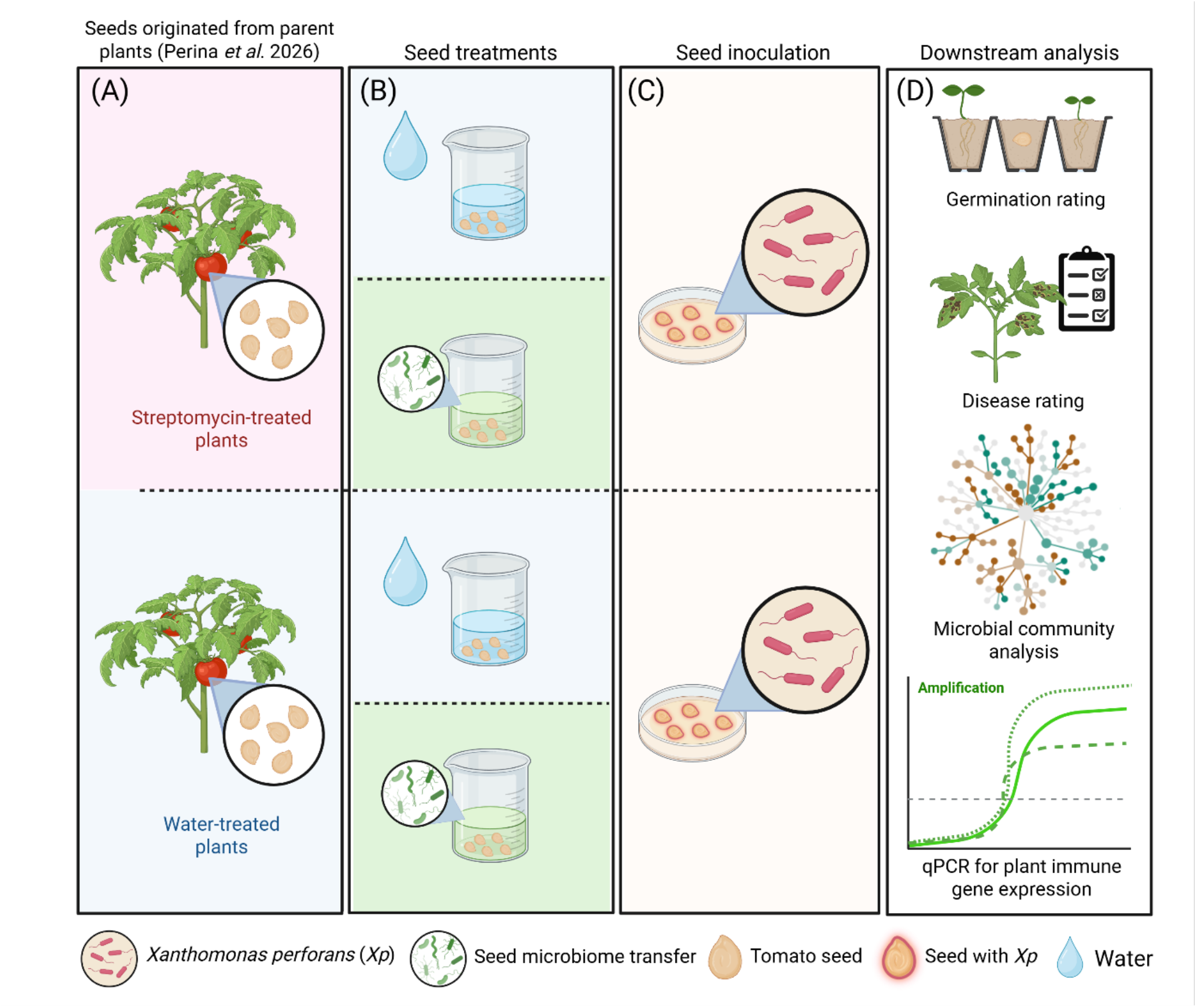
Schematic overview of the progeny and seed microbiome transfer (SMT) experiment. Step A, Progeny seeds were obtained from 27 parental plants previously treated with either water or streptomycin. Step B, Harvested seeds received either water (control) or the SMT inoculant. Step C, all seeds were challenged with the streptomycin-resistant *Xanthomonas perforans* strain Xp1-6 on LB media. Step D: Seed germination and disease severity were assessed; rhizosphere and plant samples were collected for downstream analyses, including 16S rRNA gene amplicon sequencing and qPCR analysis of plant immune gene expression, respectively.

The SMT inoculant was prepared from seeds collected from water-treated parental plants without prior pesticide exposure following a published protocol [26]. Seeds were longitudinally halved [24], incubated in Removal Buffer B (2 mL g⁻¹ seed) for 2.5 h at 20 °C and 150 rpm, and centrifuged at 10,000 × g for 10 min after removal of seed tissue. Pellets were resuspended in sterile water and adjusted to OD600 = 0.01. Seeds were pooled by parental treatment from two fruits per plant and randomly subsampled to 40 seeds per treatment. SMT seeds were incubated in the microbial suspension for 30 min at 20 °C and 70 rpm following the approach of Simonin et al. [27]; control seeds underwent the same procedure in sterile water.

Following SMT treatment, seeds were surface-dried on sterile filter paper and inoculated with *X. perforans* XP1-6 by contacting both seed surfaces with actively growing colonies on LB agar. Seeds were incubated in contact with colonies for 24 h at 25 °C and then aseptically sown.

### Growth chamber experiment

Progeny seeds (±SMT and +*X. perforans*, as described above) were sown in 900-mL pots filled with the commercial peat-perlite substrate (Jolly Gardener HFC 20). All progeny plants were cultivated in a controlled plant growth chamber under specific conditions (16 h light / 8 h dark; temperature 26°C (day), 24°C (night); humidity approximately 70%; and photosynthetic photon flux density (PPFD) 250–300 µmol m^−2^ s^−1^ at canopy level) to minimize environmental variation and standardize physiological and molecular responses, especially for RT-qPCR defense gene expression analyses. Plants were irrigated daily to field capacity.

### Seed germination and disease severity assessment

Seed germination was assessed as the percentage of seeds producing emerged seedlings. Five plants per treatment were evaluated in each of two independent experimental runs (n = 10 per treatment). Disease severity was assessed using a standardized area diagram scale for bacterial spot of tomato, developed and validated by Duan et al. [28], which accounts for both chlorotic and necrotic foliar symptoms. Five replicates per treatment were arranged in five blocks across two independent experimental runs (n = 10 per treatment). Disease severity was assessed beginning ∼14 days after germination, when symptoms first appeared, and every 2 days thereafter for six assessments. Plants were irrigated to field capacity and sprayed every 48 h to maintain uniform leaf wetness and disease pressure. The progression of bacterial spot disease was summarized using the area under the disease progress curve (AUDPC), calculated by Shaner & Finney’s (1977) method: AUDPC = Σ [(yi + yi+1) / 2] × (ti+1 − ti), where yi is severity at assessment i, ti is the time (days), with the sum over all intervals.

### RNA isolation and gene expression analyses by qRT-PCR

Tomato leaf samples were collected 5 days after seedling emergence and were pulverized under liquid nitrogen and total RNA was extracted by Direct-zol RNA MiniPrep Kit (Cat. R2050; Zymo Research, Irvine, CA, USA) following the manufacturer’s protocol. Total RNA was resuspended in nuclease-free water to 30-50 µl^−1^ nuclease free water. First-strand cDNA was synthesized from DNase-treated RNA using the Maxima First Strand cDNA Synthesis Kit for RT-qPCR (Cat. No. K1671; Thermo Fisher Scientific, Cleveland, OH, USA) in 20-µL reactions following the manufacturer’s instructions. Quantitative PCR reactions were performed using the Apex GREEN qPCR Master Mix (2×), without ROX dye (Cat. No. 42-116PG; Genesee Scientific, San Diego, CA, USA). Each reaction was prepared in a final volume of 20 µL on ice and contained: 10 µL Apex GREEN (2×); 0.2 µL of each primer (final concentration 0.01 mM); 6.8 µL nuclease-free water; and 3.0 µL of 1:5 diluted cDNA. Reactions were assembled in 96-well qPCR plates, with each sample run in technical triplicate. Amplification was performed on a Bio-Rad CFX Opus 96 Real-Time PCR System under the following thermal cycling conditions: initial denaturation/polymerase activation at 95°C for 15 min; 39 cycles of denaturation at 95°C for 15 s, annealing at 59°C for 30 s (with fluorescence acquisition at this step), and extension at 72°C for 30 s; followed by a melt curve analysis from 65°C to 95°C (0.5°C increments, 5 s hold) to confirm amplicon specificity. The gene-specific primers are listed in Table 1. Transcript abundance data for tomato was normalized against expression of *COX* housekeeping gene following the 2^−ΔΔCt^ method for relative quantification. Gene expression was normalized against the mean ΔCt of the progeny of water-control mother plants without SMT treatment (*Xp*+water). in CFX Maestro software (Bio-Rad) with automated baseline and threshold adjustments to extract Ct values. Normalized expression values (2^−ΔΔCt^, relative to the reference gene *COX*) was analyzed for statistical differences for three defense genes (*CHI14*, *Flg_Sens*, and *Gluc*). Between-treatment differences were assessed for each gene using linear mixed-effects models with treatment as a fixed effect and experiment and block-within-experiment as random effects, followed by Tukey-adjusted pairwise comparisons (nlme package and emmeans packages)[29, 30]. The relationship between defense gene expression and disease severity was evaluated by multiple regression (AUDPC ∼ *CHI14* + *Flg_Sens* + *Gluc*) with multivariate multiple regression treating the three genes as a joint response (Wilks’ lambda).

**Table 1.**
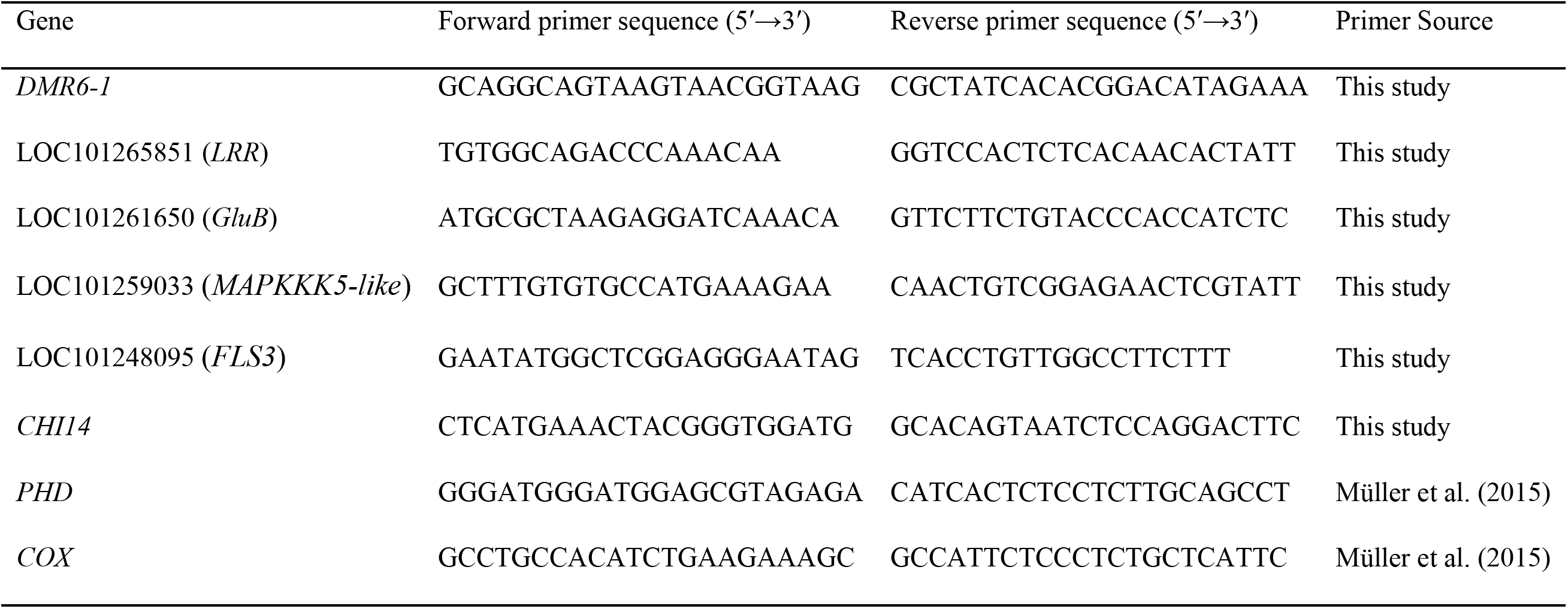
Primer sequences designed and optimized for RT-qPCR analysis of tomato defense/ISR gene expression. *DMR6-1*, Downy Mildew Resistance 6-like 1, a detoxification enzyme potentially affected by antibiotic treatment; *LRR*, leucine-rich repeat protein 1-like, involved in pathogen recognition and defense signaling; *Gluc*, β-1,3-glucanase, involved in pathogen cell-wall degradation and defense responses; *MAPKKK5-like*, mitogen-activated protein kinase 5-like, involved in defense signaling; *FLS3*, flagellin sensing 3, involved in recognition of *Xanthomonas perforans* flagellin; *CHI14*, chitinase 14, involved in pathogen cell-wall degradation and defense signaling. *PHD* was used as the reference gene for normalization, and *COX* was included as an additional control gene [31].

### Physiological statistical approach

Germination percentage and AUDPC were analyzed by one-way ANOVA followed by Tukey’s HSD test for pairwise comparisons among treatments R version 4.6.1 [30], using rstatix package [32]. When applicable, compact letter displays were generated using multcompView [33]. Germination percentage and AUDPC were visualized using boxplots generated with ggplot2 [34]. Normalized RT-qPCR expression values were averaged across technical replicates before analysis. Associations between AUDPC and immune-associated gene expression (*CHI14*, *Flg_Sens*, and *GluB*) were assessed using Spearman’s rank correlation. Linear regression lines with 95% confidence intervals were included for visualization only.

### Microbiome rhizosphere sampling

At 24 days after inoculation (DAI), rhizosphere and ripening-stage fruit samples from experiment 1 were collected for seed microbiome analysis and stored at −80 °C until DNA extraction. Approximately 7 g of root material with adhering soil was aseptically transferred to 50 mL tubes containing 25 mL potassium phosphate buffer (6.75 g KH₂PO₄, 8.75 g K₂HPO₄, 0.01% Triton X-100, pH 7.4, per L). Samples were kept on ice, vortexed for 30 s, and ultrasonicated for 10 min at ambient temperature (Vevor Ultrasonic Cleaner, Guangdong, China). Roots were removed, and suspensions were centrifuged at 4,000 × g for 10 min at 4 °C. The supernatant was discarded, and the rhizosphere microbial pellet was stored at −80 °C until DNA extraction.

### DNA extraction, quality control, sequencing and microbiome bioinformatic analysis

Genomic DNA was extracted from rhizosphere samples using the Zymo Quick DNA Faecal/Soil Microbe MiniPrep Kit (Zymo Research, USA) according to the manufacturer’s instructions. DNA concentration and quality were assessed using a NanoDrop spectrophotometer (Thermo Fisher).

Rhizosphere DNA samples (n = 40) were sequenced by SeqCenter (Pittsburgh, PA, USA) targeting the V3-V4 regions of 16S rRNA using a MiSeq 622-cycle flow cell, producing 2 × 301 bp paired-end reads.

Amplicon sequence data were processed using Mothur following the MiSeq SOP [35]. OTU, taxonomy, phylogenetic, and metadata tables were imported into phyloseq [36]. Samples were rarefied to the minimum library size using rarefy_even_depth. Beta diversity was assessed using unweighted UniFrac distances, and treatment effects were evaluated by PERMANOVA with 999 permutations using adonis2 in vegan [37]. Centroid distances were compared between treatments using one-way ANOVA and Tukey’s HSD post hoc tests with vegan, dplyr, and ggplot2 [34]. Phylogenetic turnover among microbial communities was quantified using the *β*-nearest taxon index (*β*-NTI) calculated from CSS-normalized OTU tables. *β*-NTI values were generated using the microecopackage [38]. Thresholds of |*β*-NTI| = 2 were used to distinguish deterministic from stochastic community assembly processes.

Spearman correlations assessed associations between AUDPC, PC2 from unweighted UniFrac ordinations, and genus-level relative abundances. P-values were adjusted using the Benjamini– Hochberg procedure; as no associations remained significant after FDR correction, nominal significance (*p* < 0.05) was reported. LDA-style scores estimated treatment-associated taxon enrichment and were visualized using ggplot2. Shared and unique OTUs among treatments were visualized using an UpSet plot generated with the UpSetR package in R [39]. OTU abundances were converted to presence–absence, and taxa detected in ≥3 samples per treatment were retained. Treatment-specific co-occurrence networks were constructed from filtered OTU tables using significant Spearman correlations (*p* < 0.05) and optimized for sparsity. Networks were visualized and explored using Gephi v0.10.1[40].

## Results

### Plant Physiological Responses to Streptomycin and Seed Microbiome Transfer

The following study examined the effects of Streptomycin (Strep) application and seed microbiome transfer (SMT) on tomato (*Solanum lycopersicum*) rhizosphere bacterial community composition, plant immune gene expression of selected genes, germination, and bacterial spot disease severity caused by *Xanthomonas perforans* (*Xp*). Germination success varied notably across treatments, as shown in Figure 2. *Xp*+Water showed the highest germination rate (98.0 ± 6.3%), followed by *Xp* +SMT (94.0 ± 8.4%). Both Streptomycin-treated groups (*Xp*+Strep) and *Xp*+Strep+SMT) had significantly reduced germination rates compared to the controls (*p = 0.015 and p = 0.002*), with no difference between them, indicating that microbiome transfer did not restore germination success under antibiotic treatment. Disease severity differed significantly among all four treatments (*p* < 0.01). *Xp*+SMT had the lowest AUDPC (45.7 ± 9.4), indicating the strongest disease suppression. *Xp*+Water showed intermediate severity (147.7 ± 7.7), while *Xp*+Strep+SMT (109.9 ± 12.1) was significantly lower than *Xp*+Strep (190.0 ± 12.0), which had the highest disease severity. Overall, parental streptomycin treatment increased disease severity in the progeny, whereas SMT reduced disease severity even following parental antibiotic exposure.

**Figure 2.**
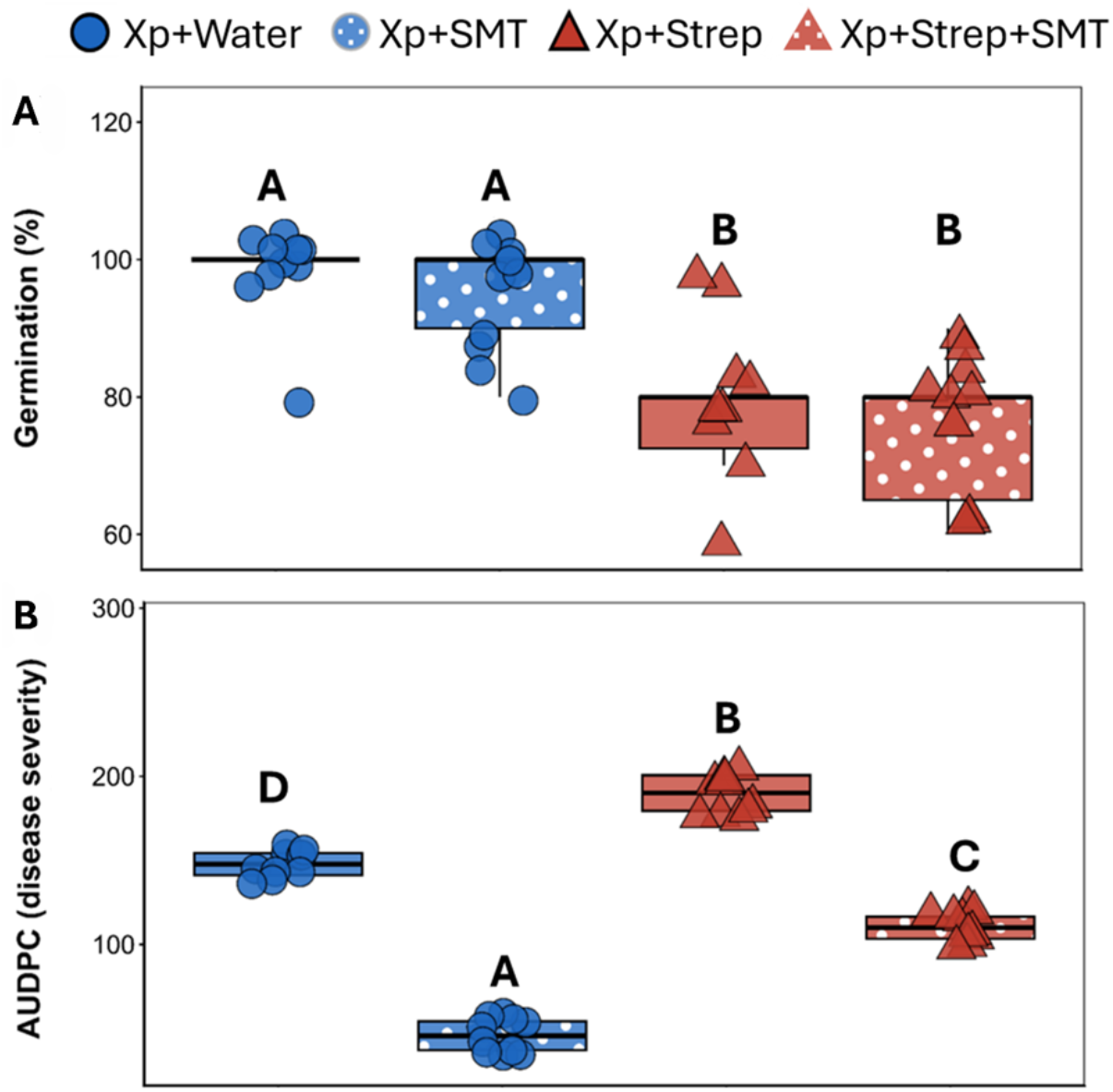
(A) Germination rate and (B) disease severity (AUDPC) were analyzed across different treatments using one-way ANOVA and Tukey HSD. Different letters above the boxplots indicate significant differences between the treatments (*p* < 0.01).

### Association between community structure and disease severity

Order-level taxonomic differences were visualized using differential heat trees across three pairwise comparisons (Figure 3). Relative to *Xp*, *Xp*+SMT enriched 127 bacterial orders versus 104 in *Xp* (Figure 3A), while *Xp*+Strep showed the largest shift, with 150 enriched orders versus 85 in *Xp* (Figure 3B). Within the streptomycin background, enrichment was more evenly distributed between *Xp*+Strep+SMT and *Xp*+Strep (120 vs. 126 orders; Figure 3C). These patterns indicate that parental streptomycin-induced dysbiosis altered progeny rhizosphere community composition, while SMT shifted the distribution of enriched taxa rather than increasing overall taxonomic richness.

**Figure 3.**
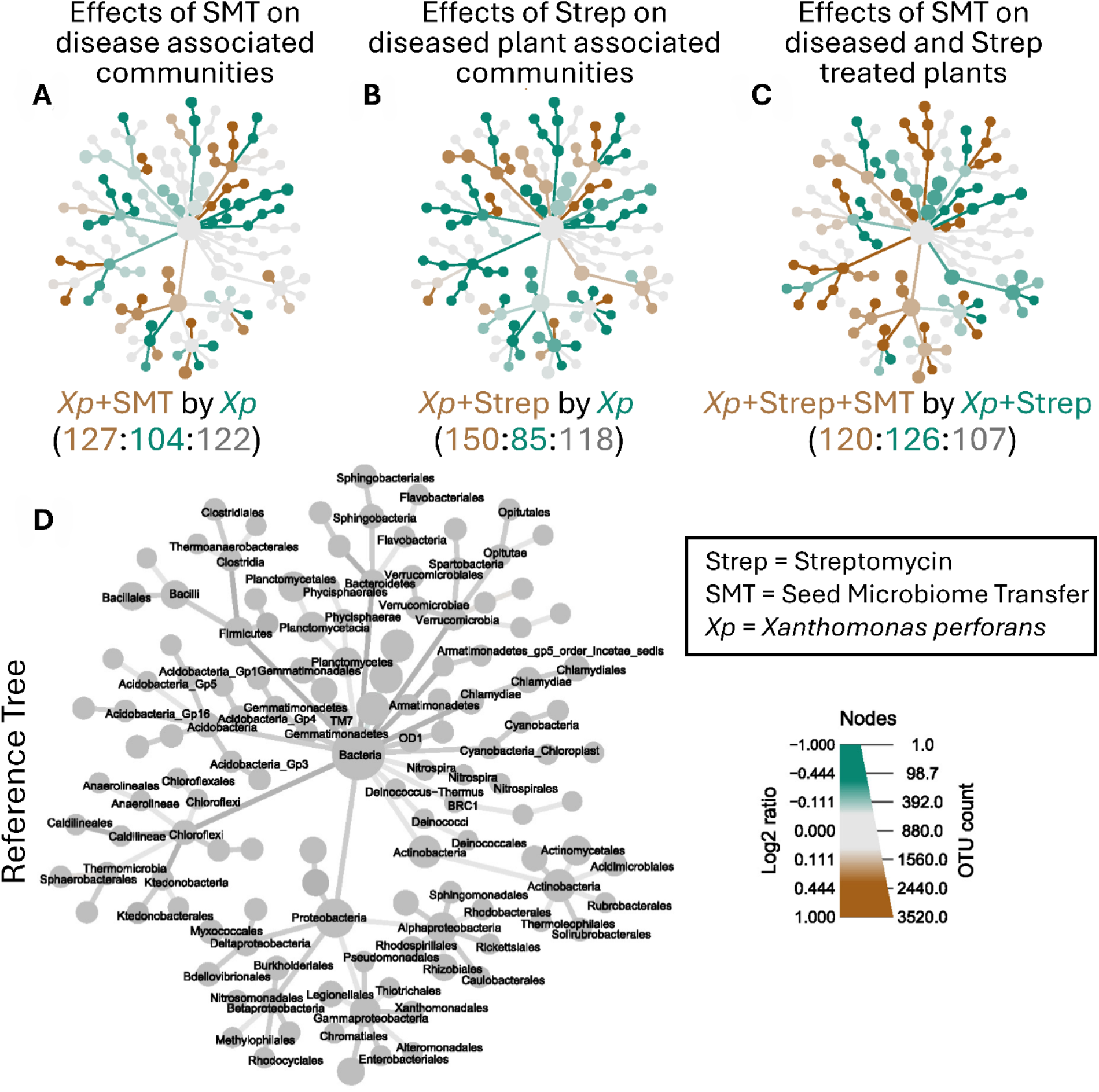
Differential heat trees showing order-level bacterial enrichment in progeny rhizosphere communities following parental streptomycin exposure, *Xanthomonas perforans* (*Xp*) challenge, and seed microbiome transfer (SMT). (A) Comparison of *Xp*+SMT and *Xp,* showing 127 orders enriched in *Xp*+SMT, 104 enriched in *Xp*, and 122 with no differential enrichment. (B) Comparison of *Xp*+Strep and, showing 150 orders enriched in *Xp*+Strep, 85 enriched in *Xp*, and 118 with no differential enrichment. (C) Comparison of *Xp*+Strep+SMT and *Xp*+Strep, showing 120 orders enriched in *Xp*+Strep+SMT, 126 enriched in *Xp*+Strep, and 107 with no differential enrichment. (D) Reference tree showing the complete order-level bacterial taxonomy. Node color indicates the direction of enrichment (brown, first-named treatment; teal, second-named treatment; gray, no differential enrichment). Node size is proportional to the total OTU abundance assigned to each taxon.

Rhizosphere community composition differed significantly among treatments (unweighted UniFrac PERMANOVA, p = 0.05), indicating shifts driven mainly by rare phylogenetic lineages (Figure 4). PCoA showed that *Xp*+Strep+SMT communities were the most dispersed, while *Xp*+Water clustered most tightly, suggesting greater stability in the absence of Streptomycin and SMT. Distance-to-centroid analysis confirmed significant differences in dispersion (ANOVA, p < 0.001), with *Xp*+Strep+SMT showing the highest variability (0.249 ± 0.008). *β*-NTI values for all samples did not fall within the stochastic range (−2 < *β*-NTI < 2), indicating that rhizosphere community assembly was governed predominantly by deterministic selection, and did not differ significantly among treatments.

**Figure 4.**
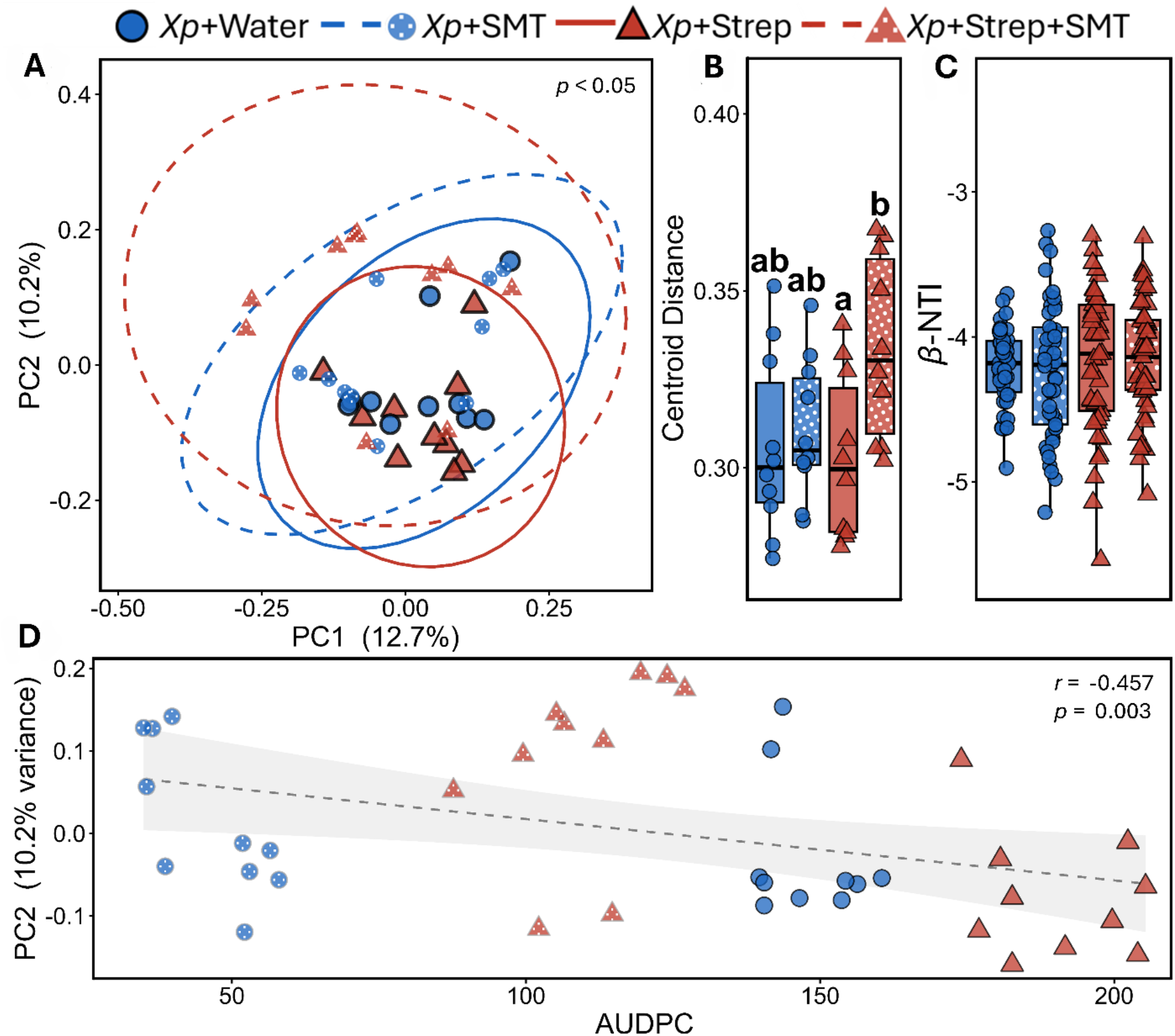
(A) Principal coordinates analysis (PCoA) of unweighted UniFrac distances showing rhizosphere bacterial community composition across four treatments. (B) Centroid distances by treatment analysis were compared with Tukey HSD, *p* < 0.05. (C) Beta-NTI values by treatment. (D) Spearman rank correlation between UniFrac PC2 scores and area under the disease progress curve (AUDPC), linking rhizosphere community structure to foliar disease severity.

To directly link rhizosphere community composition to plant disease outcomes, the second principal coordinate (PC2) from the unweighted UniFrac PCoA was correlated with AUDPC values across all 40 samples. A significant negative Spearman correlation was detected (r = −0.426, P = 0.006), indicating that samples with higher PC2 scores characteristic of *X*p +SMT communities experienced lower disease severity. In contrast samples with lower PC2 scores predominantly *Xp*+Strep+Water were associated with the highest AUDPC values. This finding establishes a statistically significant link between rhizosphere community structure and foliar bacterial spot severity in tomato. Treatment responses were consistent across experimental batches, with no apparent batch-associated differences in the measured response variables (Supplemental Figure 1-2). Treatment effects were therefore considered reproducible across batches.

UpSet analysis revealed a highly resilient core rhizosphere microbiome shared across all treatments (Figure 5A). Of the total OTUs detected, 1,834 were present in all four treatments, indicating strong stability of the dominant microbial community despite Streptomycin application and SMT. However, richness differed among treatments, with *Xp*+Strep+SMT exhibiting the greatest total richness (5,904 OTUs) and the highest number of unique OTUs (648), compared to *Xp*+Water (124), *Xp*+SMT (191), and *Xp*+Strep (173).

**Figure 5.**
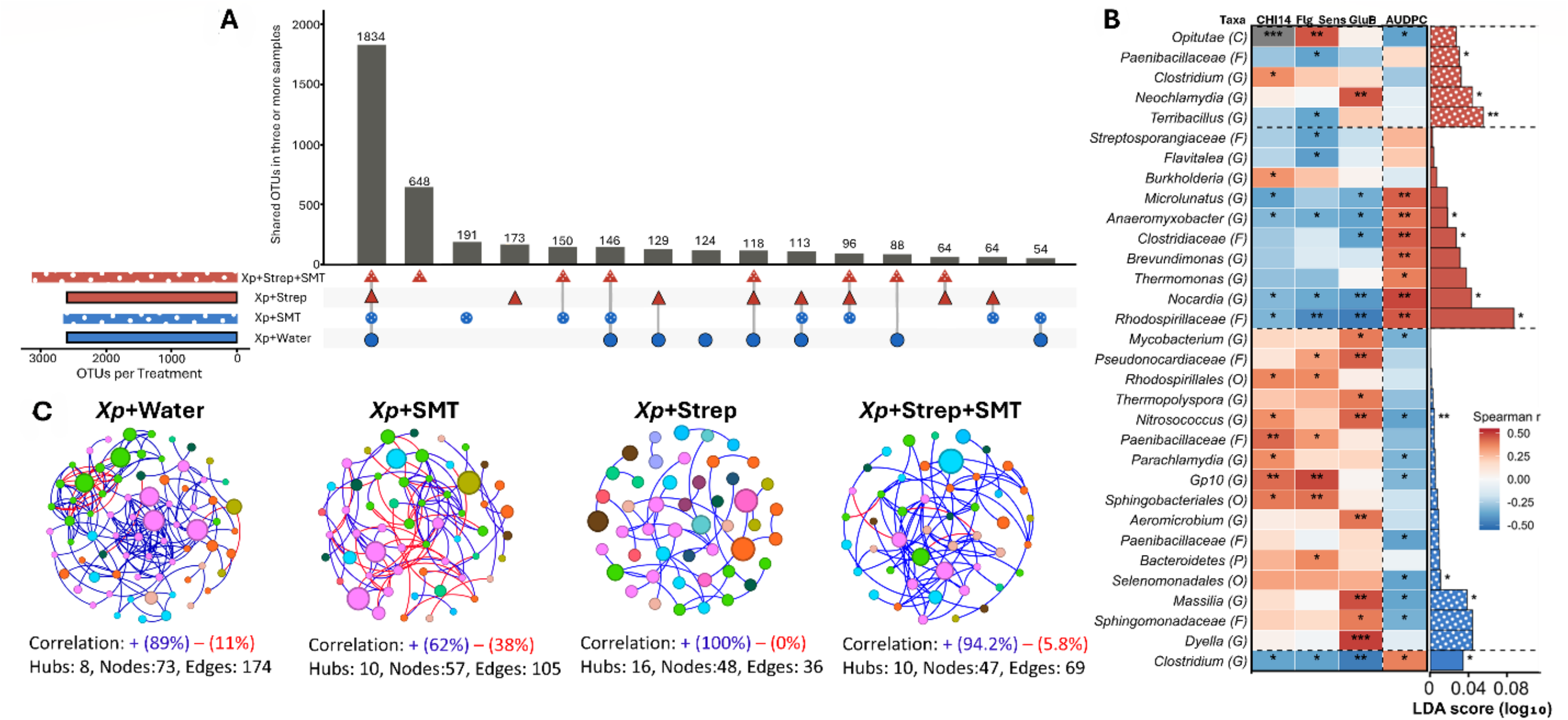
(A) UpSet plot showing the number of OTUs shared across treatment groups present in three or more samples. Horizontal bars indicate total OTU richness per treatment; vertical bars indicate the size of each intersection, with filled symbols below denoting the contributing treatments. The largest intersection (1,834 OTUs) represents OTUs shared across all four treatment groups; *Xp*+Strep+SMT harbored the greatest number of treatment-exclusive OTUs (648). (B) Heatmap of Spearman r values between genus-level relative abundance, relative plant immune gene expression (*CHI14, FLS3, GluB*) and disease severity (AUDPC), paired with LEfSe linear discriminant analysis (LDA) scores indicating treatment enrichment. Asterisks indicate significant Spearman correlations (*p* < 0.05, \**p* < 0.01, \*\**p* < 0.001). (C) Spearman rank co-occurrence networks for each treatment group, constructed from OTU relative abundances. Network statistics (percentages of positive/negative correlations, and the numbers of hubs, nodes, and edges) are shown below each network.

To identify defense genes associated with disease outcomes, Spearman correlations were calculated between the normalized expression of each candidate gene and disease severity (AUDPC) at the plant level. *CHI14*, *Flg_Sens*, and *GluB* were significantly correlated with AUDPC (p < 0.05) and were selected for subsequent analyses, whereas *DMR6*, *LRR*, and *MKinase* showed no significant association (Supplemental Figure 3). The resulting genes were compared across 369 prevalence-filtered taxa, this analysis yielded 102 nominally significant associations (*p* < 0.05) across the four response variables. *Nocardia* showed the strongest positive association with AUDPC (r = 0.491, *p* = 0.001) and was significantly negatively correlated with *CHI14* (r = −0.324, *p* = 0.042), *FLS3* (r = −0.360, *p* = 0.023), and *GluB* expression (r = −0.398, *p* = 0.011). *Nocardia* was also significantly enriched in the *Xp*+Strep+Water treatment (*p* = 0.035). *Rhodospirillaceae* (two OTU clusters) showed consistent negative associations with *GluB* (r = −0.502 to −0.491, *p* < 0.002) and positive associations with AUDPC (r = 0.424–0.443, *p* < 0.007), and was most abundant in *Xp*+Strep+Water communities. *Anaeromyxobacter* exhibited similar patterns, with negative correlations to *CHI14* (r = −0.344, *p* = 0.030), *FLS3* (r = −0.384, *p* = 0.015), and *GluB* expression (r = −0.376, *p* = 0.017), and a positive association with AUDPC (r = 0.424, *p* = 0.007). *Clostridiaceae* was also positively associated with AUDPC (r = 0.430, *p* = 0.006) and negatively associated with *GluB* expression (r = −0.371, *p* = 0.018). Collectively, these taxa form a disease-associated cluster enriched under antibiotic-driven dysbiosis.

In contrast, several taxa were associated with reduced disease severity and enhanced immune responses. *Gp10* showed strong positive correlations with *CHI14* (r = 0.432, p = 0.005) and *FLS3* (r = 0.494, P = 0.001) expression and a negative association with AUDPC (r = −0.330, p = 0.038). *Massilia* was negatively correlated with AUDPC (r = −0.441, P = 0.004) and positively with *GluB* expression (r = 0.430, p = 0.006), and was enriched in control treatments *Xp*+SMT (H = 9.75, p = 0.021). *Dyella* showed the strongest positive association with *GluB* expression (r = 0.515, p = 0.001). *Pseudonocardiaceae* and *Sphingomonadaceae* were positively correlated with *GlucB* expression (r = 0.443, p = 0.004 and r = 0.392, p = 0.012, respectively). Both taxa also showed negative correlations with AUDPC, although this association was significant only for *Sphingomonadaceae* (r = −0.342, p = 0.031) and not for *Pseudonocardiaceae* (r = −0.232, p = 0.150). Consistent with these associations, both taxa were more abundant in *Xp*+SMT communities.

These patterns were further supported by LEfSe analysis, which identified *Nocardia* and *Rhodospirillaceae* as enriched in *Xp*+Strep, and *Massilia*, *Sphingomonadaceae*, and *Gp10* as enriched in *Xp*+Water treatments, consistent with microbial signatures associated with disease susceptibility versus immune priming.

Co-occurrence network analysis revealed treatment-specific shifts in microbial interaction structure. *Xp*+Water networks were the most connected, with 73 nodes and 174 edges and predominantly positive associations (89%). *Xp*+Strep reduced network size and connectivity to 48 nodes and 36 edges, a 34% and 79% reduction, respectively, with exclusively positive associations. In the streptomycin-history background, SMT did not recover node number (47 nodes) but increased edges from 36 to 69 and mean degree from 1.50 to 2.94, resulting in network density of 0.064, similar to *Xp*+Water (0.066). *Xp*+SMT contained 57 nodes and 105 edges and exhibited the highest proportion of negative associations (38%), compared with 11% in *Xp*+Water. Thus, parental streptomycin exposure reduced network connectivity, whereas SMT increased connectivity within the dysbiotic background and altered association structure in *Xp* plants.

### Defense gene expression is negatively correlated with disease severity and partially restored by SMT

Normalized expression of *CHI14*, *Flg_Sens*, and *GluB* differed among treatments based on Tukey-adjusted pairwise comparisons (Supplementary Figure 4). All three genes showed highest expression in *Xp*+SMT. *CHI14* expression was significantly higher in *Xp*+SMT than all other treatments, with *Xp*+Strep+SMT intermediate and higher than *Xp*+Water. *GluB* expression was higher in both SMT treatments than in *Xp*+Water or *Xp*+Strep, which did not differ. *Flg_Sens* expression was higher in *Xp*+SMT than *Xp*+Water and *Xp*+Strep, while *Xp*+Strep+SMT was intermediate. Thus, SMT increased defense gene expression, whereas streptomycin alone did not significantly alter expression relative to *Xp*+Water.

To examine the relationship between defense-associated gene expression and disease outcomes, Spearman correlations were calculated between AUDPC and the relative expression of *CHI14*, *FLS3*, and *GluB* across all treatment groups (Figure 6). Expression of each gene was negatively associated with disease severity: *CHI14* (r = −0.728, p < 0.001), *FLS3* (r = −0.634, p < 0.001), and *GluB* (r = −0.550, p < 0.001). Thus, across the complete dataset, samples with higher expression of these genes generally exhibited lower AUDPC values. However, expression varied within treatments, and the overlap among treatment groups did not support a consistent treatment-specific expression pattern based on these correlations alone. Together, these results indicate that parental streptomycin-induced dysbiosis was associated with suppressed defense gene expression in progeny plants and increased susceptibility to *X. perforans*, and that SMT partially mitigated this effect.

**Figure 6.**
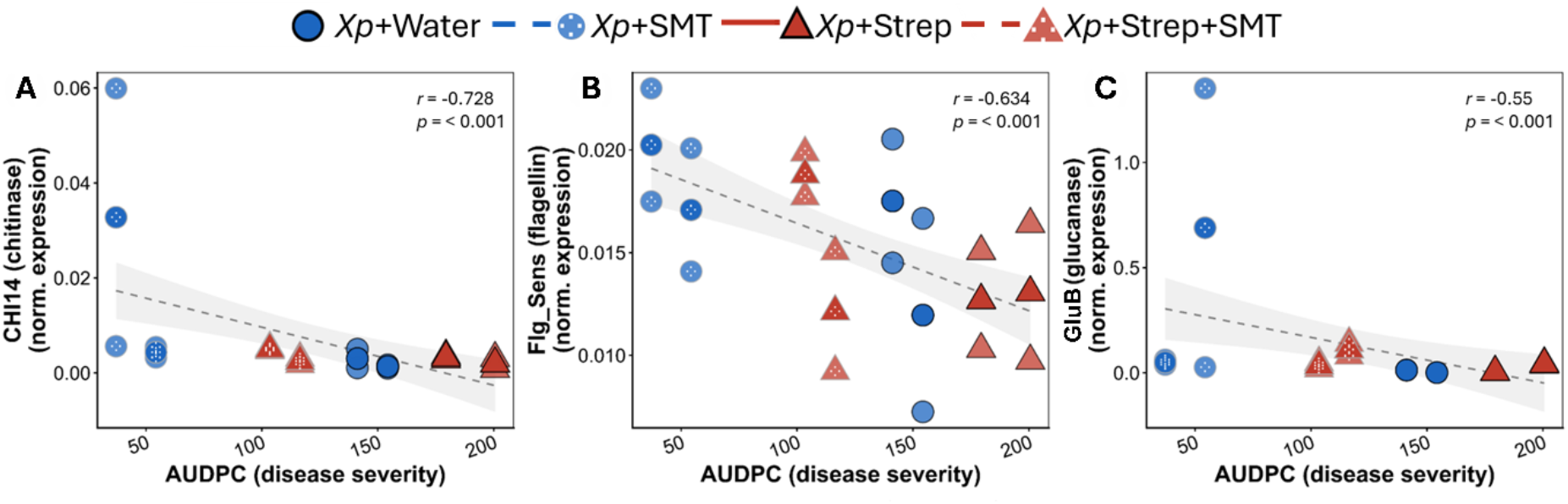
(A–C) Spearman correlations between AUDPC and normalized expression of the defense-associated genes *CHI14*, *Flg_Sens*, and *GluB*. Higher immune gene expression was significantly associated with lower disease severity (all *p* < 0.001). Dashed lines represent linear regression fits with 95% confidence intervals. Blue = control treatments; red = Streptomycin treatments (Strep); circles = control; triangles = Streptomycin; polka dot fill = Seed Microbiome Transfer (SMT).

## Discussion

This study shows that parental streptomycin exposure was associated with measurable intergenerational effects in tomato progeny, including reduced germination, altered rhizosphere bacterial community composition, and increased susceptibility to *Xanthomonas perforans*. Across treatments, expression of the defense-associated genes *CHI14*, *Flg_Sens,* and *GluB* was negatively correlated with disease severity. However, these associations alone do not demonstrate treatment-induced suppression of defense gene expression. Although the effects of host stress on microbiome dynamics have been increasingly studied, whether these effects persist across generations remains underexplored [24]. Here, we show that seed microbiome transfer (SMT) from healthy plants donors partially reversed these intergenerational effects, restoring aspects of rhizosphere community structure, recovering defense gene expression, and significantly reducing foliar disease severity. The strong negative correlations between *CHI14, Flg_Sens,* and *GluB* expression and disease severity provide a mechanistic link between microbiome disruption, impaired immune priming, and increased disease susceptibility, positioning the seed microbiome as a functional bridge between parental dysbiosis and progeny plant health.

The intergenerational effects of dysbiosis observed here extend evidence that rhizosphere microbial legacies can influence plant health across generations [23]. In rice, rhizosphere communities associated with leaf disease generated persistent changes in progeny microbiome composition and disease outcomes [23]. Similarly, parental streptomycin exposure produced persistent compositional shifts in progeny rhizosphere communities, indicating that microbiome disruption can extend beyond the directly exposed generation. These communities showed pronounced restructuring and depletion of abundant, taxonomically diverse bacterial orders, consistent with the broad-spectrum activity of streptomycin and selection for antibiotic-tolerant taxa [6, 7]. This restructuring is also consistent with the Anna Karenina Principle of plant dysbiosis, whereby host stress is associated with increased microbiome variability and reduced community stability [14]. Together, these findings indicate that antibiotic-induced dysbiosis can generate intergenerational microbial legacies, potentially mediated by seed-associated microbial transmission, that alter progeny rhizosphere assembly and plant–microbe interactions [24, 26].

Seeds from streptomycin-treated parents exhibited reduced germination (*p* = 0.015–0.002), indicating an intergenerational effect of parental microbiome disruption. SMT did not restore this phenotype, contrasting with its effects on disease severity and suggesting that early developmental traits may depend on seed-intrinsic microbial communities, particularly endophytes within the seed endosphere, that are not readily re-established by external inoculation. Endophytes colonize internal seed tissues during fruit development through floral and vascular pathways and are therefore largely inaccessible to surface-based inoculation [3, 26]. Parental streptomycin exposure may consequently disrupt the transmission or functional integrity of core endophytic taxa involved in seedling establishment and early development. Restoration strategies targeting the seed endosphere, including in planta interventions during parental development, may therefore be required to recover developmental traits affected by parental antibiotic exposure.

A key finding of this study was that *Xp*+SMT plants exhibited the lowest disease severity among all treatment groups, representing a substantial reduction compared with *Xp*+Water controls. This protection may involve two non-mutually exclusive mechanisms. First, SMT may have introduced microbial functions that modulated host defenses through microbiome-mediated induced systemic resistance (ISR) [1, 24], consistent with disease suppression by synthetic microbial communities in tomato and other crops [34, 35]. Second, because SMT was applied directly to seeds before pathogen inoculation, transferred microorganisms may have restricted *X. perforans* establishment through priority effects, competition for space or nutrients, or direct antagonism on the seed surface [41]. Because our experimental design did not independently quantify pathogen establishment on the seed surface and the contribution of host immune responses, the relative importance of these mechanisms could not be distinguished. Within the context of antibiotic-induced dysbiosis, SMT partially alleviated the elevated disease susceptibility associated with parental streptomycin exposure, as evidenced by reduced disease progression in *Xp*+Strep+SMT plants compared with *Xp*+Strep plants. These findings suggest that SMT can recover disease-suppressive functions without fully re-establishing the original microbial community. Reintroduction of naturally occurring epiphytic microbiota onto tomato seeds has similarly been shown to reduce pathogen establishment and disease in seedlings [42]. More recently, seed-associated bacteria were shown to competitively exclude *Xanthomonas campestris* through siderophore-mediated iron competition at the seed–microbiome interface[41].Thus, functional replacement and/or competition with *X. perforans* during seed colonization may have contributed to the observed disease suppression, although these mechanisms were not directly evaluated in our study.

Genus-level analyses linked intergenerational antibiotic-induced community restructuring with host immune suppression, revealing microbial signatures associated with disease susceptibility and resistance. A disease-associated cluster comprising *Nocardia*, *Rhodospirillaceae*, *Anaeromyxobacter*, and *Clostridiaceae* was enriched under streptomycin-induced dysbiosis and positively associated with AUDPC but negatively with defense gene expression, consistent with proliferation following antibiotic-mediated depletion of beneficial competitors [7, 9, 15]. In contrast, *Massilia*, *Dyella*, Acidobacteria subgroup 10 (Gp10), and *Sphingomonadaceae* were enriched in control and SMT treatments and positively associated with defense gene expression but negatively with disease severity, consistent with their reported roles in plant growth promotion, immune priming, and pathogen suppression [1, 15, 22]. LEfSe analysis further supported these taxa as candidate biomarkers of disease-suppressive rhizosphere states [15, 19]. The coordinated suppression of *CHI14*, *Flg_Sens*, and *GluB* in progeny of streptomycin-treated plants, together with their negative associations with disease severity, suggests reduced pattern-triggered immunity. Their partial recovery following SMT further supports microbiome-mediated restoration of immune function across generations, consistent with the extended plant immune system concept [1, 22].

The intergenerational consequences of streptomycin-induced dysbiosis demonstrated here highlight a largely overlooked cost of antibiotic use in agriculture: the disruption of plant-associated microbial communities does not end with the treated generation but may propagate through the seed microbiome, compromising the immunity and disease resistance of progeny plants. Streptomycin, one of the most widely used antibiotics for control of bacterial plant diseases [6, 11], can exert effects that extend well beyond the treated generation, generating intergenerational microbiome legacies with measurable consequences for progeny immunity and disease susceptibility. In this study, parental streptomycin exposure was associated with persistent changes in progeny microbiome composition, immunity, and disease susceptibility. However, our experimental design did not determine whether these effects were transmitted directly through the seed microbiome. Streptomycin-induced changes in other aspects of seed biology, such as seed physiology, chemistry, or immune status, may also have influenced subsequent microbiome recruitment. Nevertheless, the persistence of these effects in the progeny under the conditions tested is consistent with rhizosphere legacy effects reported in rice and seed microbiome transmission studies in legumes [25, 26]. However, our results demonstrate that deliberate SMT from healthy plant donors represents a practical, biology-based strategy to partially mitigate these legacy effects, extending the “patching the leak” framework to an intergenerational dimension and contributing to the growing evidence that microbiome-based interventions can restore holobiont health disrupted by external stressors [19, 43]. Future work should identify the specific taxa and functional traits within the SMT inoculant responsible for protection and distinguish host-mediated ISR from direct microbial antagonism by evaluating SMT effects with and without pathogen inoculation. It should also determine whether protection persists across successive generations and whether enriching donor inocula with beneficial taxa, such as *Massilia* or members of *Sphingomonadaceae,* enhances efficacy. Particular attention should be given to *Sphingomonas*, a recurrent tomato phyllosphere colonizer that can include streptomycin-resistant strains and has demonstrated protective activity against foliar bacterial pathogens [44–46]. Such strains may persist following antibiotic exposure and directly compete with *X. perforans* for foliar niches, a hypothesis that should be tested through strain isolation, antibiotic-susceptibility assays, and co-colonization experiments. Therefore, synthetic communities defined to incorporate these traits could provide a more reproducible and targeted alternative to whole-community SMT [47, 48].

## Acknowledgements

This work was supported by the USDA National Institute of Food and Agriculture (NIFA) project no. 2022-68015-36721, the University of Florida’s Institute of Food and Agricultural Sciences (UF/IFAS) seed grant, and by the Research Capacity Fund (Hatch) program, project award no. 7010682.

## Data Availability

The 16S rRNA gene amplicon sequencing data generated in this study have been deposited in the NCBI Sequence Read Archive under BioProject accession PRJNA1504940.

## References

1. Pieterse CM. The extended plant immune system. Molecular Plant-Microbe Interactions. 2025;38:780–95

2. Trivedi P, Leach JE, Tringe SG et al. Plant–microbiome interactions: From community assembly to plant health. Nature reviews microbiology. 2020;18:607–21

3. Bergmann GE, Lui K, Lopez C et al. Contribution of floral transmission to the assembly and health impact of bacterial communities in watermelon seeds. Phytobiomes Journal. 2026;10:165–77

4. Abdelfattah A, Tack AJ, Lobato C et al. From seed to seed: The role of microbial inheritance in the assembly of the plant microbiome. Trends in Microbiology. 2023;31:346–55

5. Trivedi P, Batista BD, Bazany KE et al. Plant–microbiome interactions under a changing world: Responses, consequences and perspectives. New Phytologist. 2022;234:1951–59

6. Batuman O, Britt-Ugartemendia K, Kunwar S et al. The use and impact of antibiotics in plant agriculture: A review. Phytopathology®. 2024;114:885–909

7. Cycoń M, Mrozik A, Piotrowska-Seget Z. Antibiotics in the soil environment— degradation and their impact on microbial activity and diversity. Frontiers in microbiology. 2019;10:338

8. Yin L, Wang X, Li Y et al. Uptake of the plant agriculture-used antibiotics oxytetracycline and streptomycin by cherry radish─ effect on plant microbiome and the potential health risk. Journal of agricultural and food chemistry. 2023;71:4561–70

9. Huang Y-H, Yang Y-J, Li J-Y et al. Root-associated bacteria strengthen their community stability against disturbance of antibiotics on structure and functions. Journal of Hazardous Materials. 2024;465:133317

10. Waechter C, Fehse L, Welzel M et al. Comparative analysis of full-length 16s ribosomal rna genome sequencing in human fecal samples using primer sets with different degrees of degeneracy. Front Genet. 2023;14:1213829 10.3389/fgene.2023.1213829

11. McManus PS, Stockwell VO, Sundin GW et al. Antibiotic use in plant agriculture. Annual review of phytopathology. 2002;40:443–65

12. Fang P, Elena AX, Kunath MA et al. Reduced selection for antibiotic resistance in community context is maintained despite pressure by additional antibiotics. ISME communications. 2023;3:52

13. Thomas VE, Abeysinghe G, Goetze PK, et al. The distinguished microbiome: Differentiating microbial dysbiosis and the pathobiome of banana fusarium tropical race 4. Phytobiomes Journal. 2025;9:608–19

14. Arnault G, Mony C, Vandenkoornhuyse P. Plant microbiota dysbiosis and the anna karenina principle. Trends in Plant Science. 2023;28:18–30

15. Ketehouli T, Pasche J, Buttrós VH et al. The underground world of plant disease: Rhizosphere dysbiosis reduces above-ground plant resistance to bacterial leaf spot and alters plant transcriptome. Environmental Microbiology. 2024;26:e16676

16. Perina FJ, Thomas VE, Ketehouli T et al. Antibiotics and copper drive compartment-specific dysbiosis and functional reprogramming in tomato microbiomes. bioRxiv. 2026:2026.05. 27.728321

17. Rolfe SA, Griffiths J, Ton J. Crying out for help with root exudates: Adaptive mechanisms by which stressed plants assemble health-promoting soil microbiomes. Current opinion in microbiology. 2019;49:73–82

18. Yildirim KC, Orel DC, Okyay H et al. Enhancing seed quality and plant growth through modification of seed microbiome with beneficial endophytic bacteria. Scientia Horticulturae. 2025;354:114531

19. Ketehouli T, Goss E, Perina F et al. Patching the leak or rebuilding the boat? Evaluating targeted probiotic cyanobacteria and microbiome transplants to counteract antibiotic-driven rhizosphere dysbiosis in tomato under *xanthomonas perforans* pressure. bioRxiv. 2026:10.64898/2026.05.20.726701

20. Trinh J, Li T, Franco JY et al. Variation in microbial feature perception in the rutaceae family with immune receptor conservation in citrus. Plant Physiology. 2023;193:689–707

21. Couto D, Zipfel C. Regulation of pattern recognition receptor signalling in plants. Nature Reviews Immunology. 2016;16:537–52

22. Du Y, Han X, Tsuda K. Microbiome-mediated plant disease resistance: Recent advances and future directions. Journal of General Plant Pathology. 2025;91:1–17

23. Jobert L, Vicheth V, Czernic P et al. Rhizosphere legacy of leaf-diseased rice and its impact on next generation. Frontiers in Microbiology. 2025;16:1677271

24. Sulesky-Grieb A, Simonin M, Bintarti AF et al. Stable, multigenerational transmission of the bean seed microbiome despite abiotic stress. Msystems. 2024;9:e00951–24

25. Maillet L, Norest M, Kautsky A et al. Plant genetic bases associated with microbiota descriptors shed light into a novel holobiont generalist genes theory. Environmental Microbiology. 2025;27:e70108

26. Chesneau G, Laroche B, Préveaux A et al. Single seed microbiota: Assembly and transmission from parent plant to seedling. MBio. 2022;13:e01648–22

27. Simonin M, Préveaux A, Marais C, et al. Transmission of synthetic seed bacterial communities to radish seedlings: Impact on microbiota assembly and plant phenotype. Peer Community Journal. 2023;3

28. Duan J, Zhao B, Wang Y et al. Development and validation of a standard area diagram set to aid estimation of bacterial spot severity on tomato leaves. European journal of plant pathology. 2015;142:665–75

29. Pinheiro J, Bates D, DebRoy S et al. Package ‘nlme’. Linear and nonlinear mixed effects models, version. 2017;3:274

30. Lenth R, Singmann H, Love J et al. Package ‘emmeans’. R package version. 2019;1

31. Müller OA, Grau J, Thieme S et al. Genome-wide identification and validation of reference genes in infected tomato leaves for quantitative rt-pcr analyses. PloS one. 2015;10:e0136499

32. Kassambara A. Rstatix: Pipe-friendly framework for basic statistical tests. CRAN: Contributed packages. 2019

33. Bretz F, Hothorn T, Westfall P. Multiple comparisons using r: Chapman and Hall/CRC, 2016.

34. Wickham H Programming with ggplot2. Ggplot2: Elegant graphics for data analysis, Springer. 241–53

35. Schloss PD. Reintroducing mothur: 10 years later. Applied and environmental microbiology. 2020;86:e02343–19

36. McMurdie PJ, Holmes S. Phyloseq: An r package for reproducible interactive analysis and graphics of microbiome census data. PloS one. 2013;8:e61217

37. Dixon P. Vegan, a package of r functions for community ecology. Journal of vegetation science. 2003;14:927–30

38. Liu C, Cui Y, Li X et al. Microeco: An r package for data mining in microbial community ecology. FEMS microbiology ecology. 2021;97:fiaa255

39. Conway JR, Lex A, Gehlenborg N. Upsetr: An r package for the visualization of intersecting sets and their properties. Bioinformatics. 2017;33:2938–40

40. Bastian M, Heymann S, Jacomy M Gephi: An open source software for exploring and manipulating networks. Proceedings of the international AAAI conference on web and social media. 361–62.

41. Chesneau G, Noel A, Bréard D et al. Lactuchelins represent lipopeptide siderophores produced by pseudomonas lactucae that inhibit xanthomonas campestris. The ISME Journal. 2026;20:wrag003

42. Morella NM, Zhang X, Koskella B. Tomato seed-associated bacteria confer protection of seedlings against foliar disease caused by pseudomonas syringae. Phytobiomes Journal. 2019;3:177–90

43. Francomano E, Aci MM, Mosca S et al. Plant health in the era of global changes, holobiont biology, and microbiome-based solutions. Horticulture Research. 2026;13:uhaf364

44. Innerebner G, Knief C, Vorholt JA. Protection of arabidopsis thaliana against leaf-pathogenic pseudomonas syringae by sphingomonas strains in a controlled model system. Applied and environmental microbiology. 2011;77:3202–10

45. Vanbroekhoven K, Ryngaert A, Bastiaens L et al. Streptomycin as a selective agent to facilitate recovery and isolation of introduced and indigenous sphingomonas from environmental samples. Environmental microbiology. 2004;6:1123–36

46. Runge P, Ventura F, Kemen E et al. Distinct phyllosphere microbiome of wild tomato species in central peru upon dysbiosis. Microb Ecol. 2023;85:168–83 10.1007/s00248-021-01947-w

47. Zhou X, Wang J, Liu F et al. Cross-kingdom synthetic microbiota supports tomato suppression of fusarium wilt disease. Nature communications. 2022;13:7890

48. Schmitz L, Yan Z, Schneijderberg M et al. Synthetic bacterial community derived from a desert rhizosphere confers salt stress resilience to tomato in the presence of a soil microbiome. The ISME Journal. 2022;16:1907–20

