## Supplementary figures for "Seed Microbiome Transfer Mitigates Intergenerational Dysbiosis, Modulates Plant Defenses and Suppresses Foliar Disease"

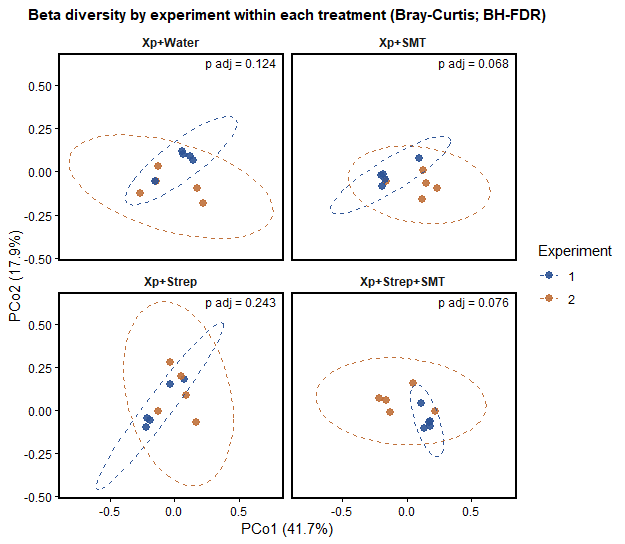

Supplementary Figure 1. **Comparison of microbial community composition between experimental batches within each treatment.** Beta diversity was assessed using Bray–Curtis dissimilarity and visualized by principal coordinates analysis (PCoA). Samples are shown by experimental batch within each treatment, with ellipses representing the dispersion of each experimental batch. PERMANOVA was used to test for differences between experimental batches within each treatment, and *P* values were adjusted using the Benjamini–Hochberg false discovery rate (BH-FDR) correction. No significant differences in microbial community composition were detected between experimental batches within any treatment after BH-FDR correction.

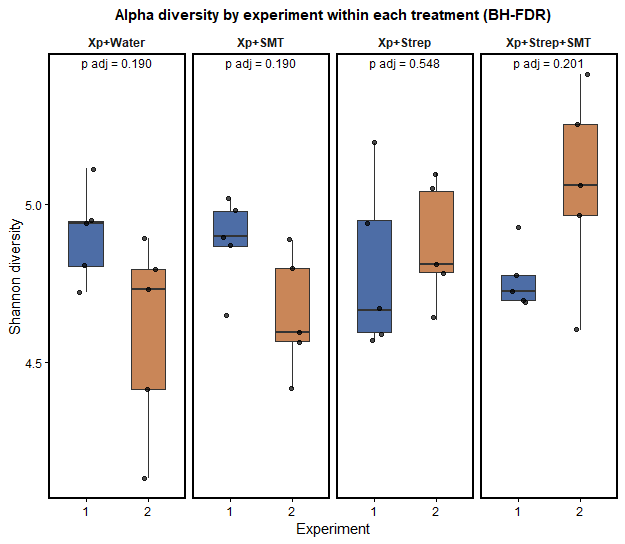

**Supplementary Figure 2. Comparison of alpha diversity between experimental batches within each treatment.** Shannon alpha diversity was calculated and visualized as boxplots for each experimental batch within each treatment. Statistical comparisons between experimental batches were performed within each treatment, with *P* values adjusted using the Benjamini–Hochberg false discovery rate (BH-FDR) correction. No significant differences in Shannon alpha diversity were detected between experimental batches within any treatment after BH-FDR correction.

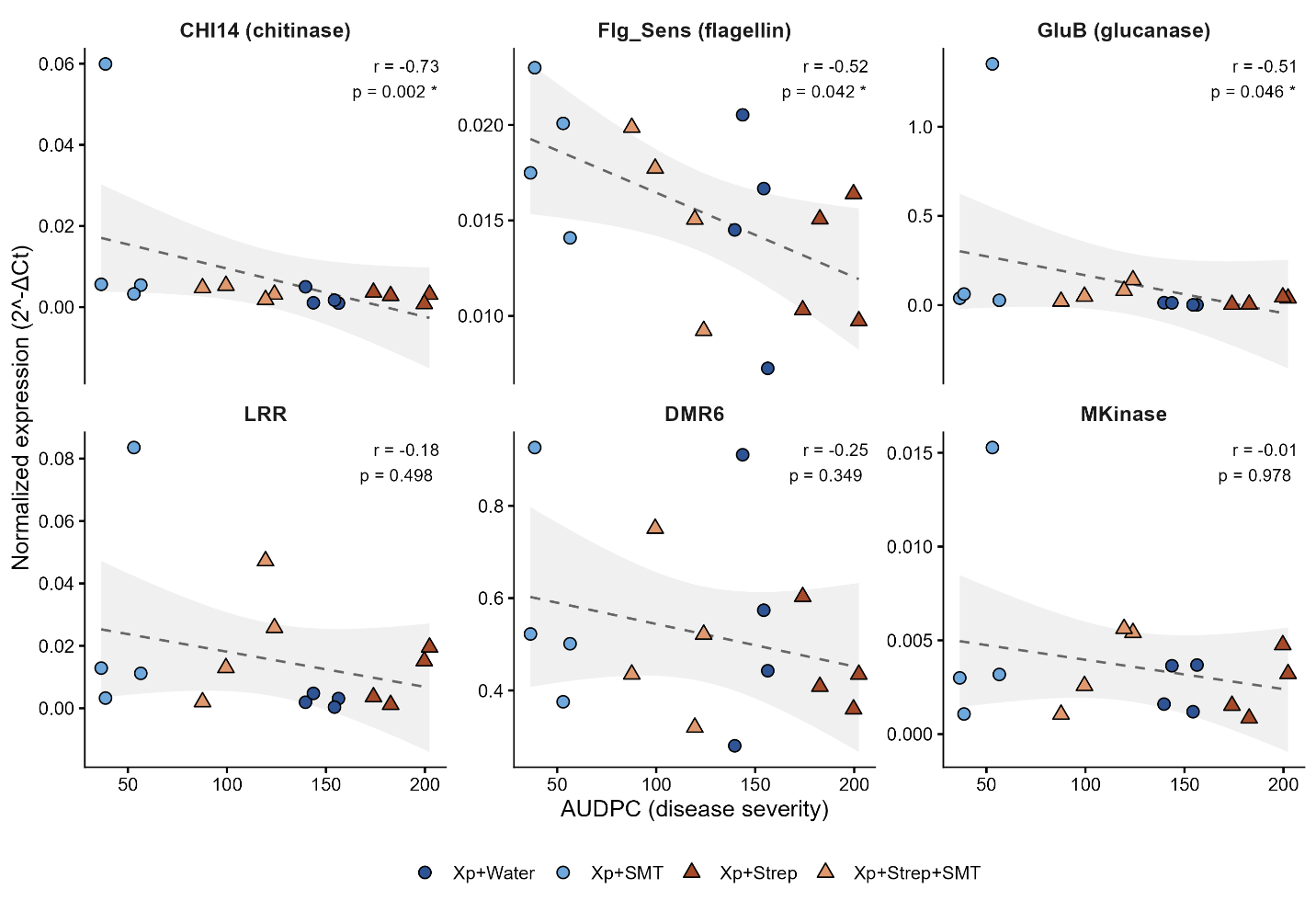

Supplemental Figure 3. **Spearman rank correlations between normalized expression of six defense-associated genes and disease severity (AUDPC) across all treatments.** *CHI14*, *Flg_Sens*, and *GluB* showed significant associations with disease severity (*p* < 0.05) and were selected for subsequent analyses, whereas *DMR6*, *LRR*, and *MKinase* showed no significant associations and were not included in downstream analyses.

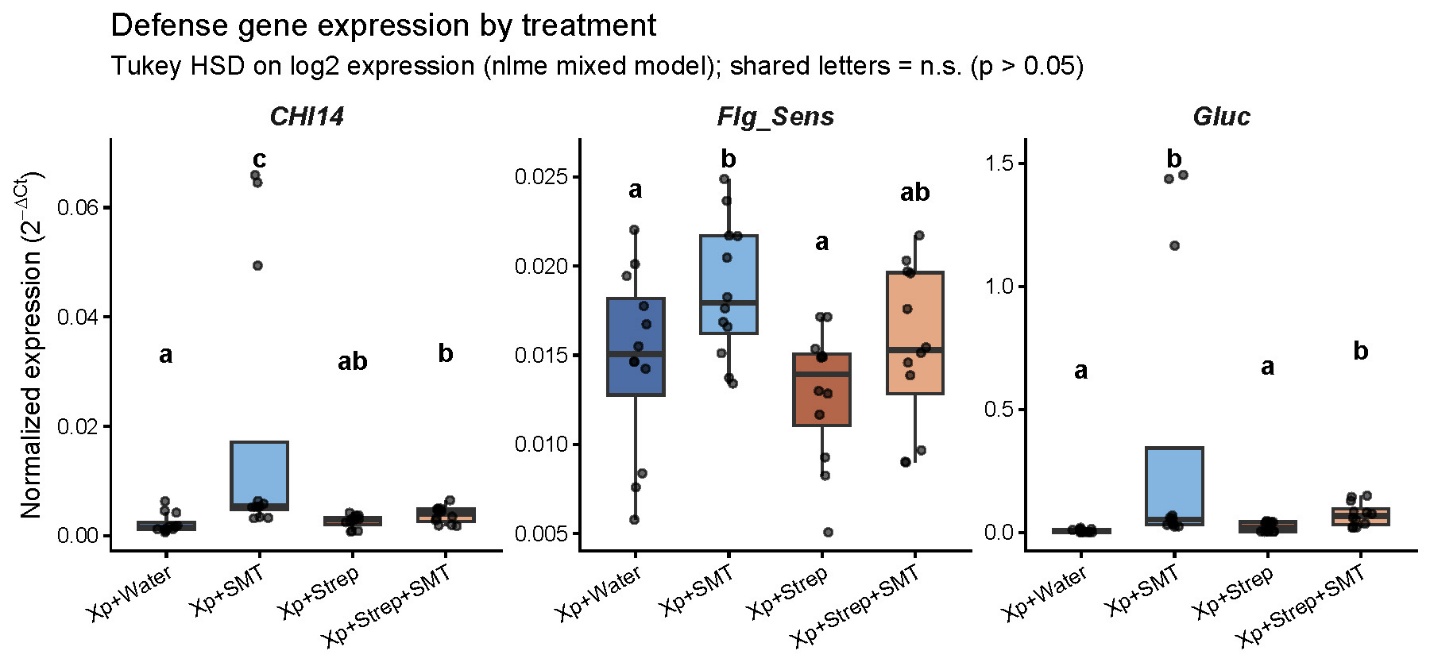

**Supplemental Figure 4.** Normalized expression of three focal defense genes (*CHI14*, *Flg_Sens*, and *Gluc*) across the four treatments (Xp+Water, Xp+SMT, Xp+Strep, Xp+Strep+SMT). Expression values (2^−ΔΔCt^, normalized to the reference gene *COX*) differences among treatments by Tukey-adjusted pairwise comparisons. Treatments sharing a letter within a panel do not differ significantly (Tukey-adjusted *p* > 0.05).
